# Towards a Rosetta Stone to Decipher the Survival Networks of Senescence

**DOI:** 10.1101/2024.07.13.603401

**Authors:** Samael Olascoaga, Norma Edith López-Diazguerrero

## Abstract

Cellular senescence, an irreversible state of cell cycle arrest in response to various stressors that can damage cells, is a key player in developing and progressing multiple chronic degenerative diseases associated with aging. Over the years, these senescent cells accumulate in organs and tissues, it is thought that the accumulation of these cells results from their capacity to evade programmed cell death by developing and activating Senescent Cell Anti-apoptotic Pathways (SCAPs); however, numerous aspects of the development and activation of these SCAPs are still unknown. In this study, we analyzed the variations in the expression levels and co-expression patterns of 39 SCAPs genes across 33 tissues from young and elderly individuals. Surprisingly, we did not observe a consistent increase in SCAPs gene expression with age in any tissue. Instead, we found tissue-specific variations in gene expression levels, changes in the strength of gene coordination, and the wiring and rewiring of gene co-expression networks that depend on the tissue. Our results suggest that the development and activation of SCAPs are far more complex than previously understood; merely increasing the expression of specific survival genes would not adequately explain the anti-apoptotic capabilities of senescent cells. We suggest that the formation and triggering of SCAPs entail a complex interplay of factors, encompassing distinct alterations in expression levels alongside shifts in the intensity and configurations of gene interactions, moreover, these modifications are unique to individual tissues. Our results deepen our understanding of how senescent cells evade programmed cell death and are essential for developing targeted pharmacological therapies that can selectively and safely eliminate senescent cells more effectively.

## Introduction

Cellular senescence is an irreversible state of cell cycle arrest triggered by various stressors, including telomere shortening, ionizing radiation, chemotherapy, activation of certain oncogenes, and oxidative stress, among others [1]. Senescent cells (SCs) exhibit a series of characteristic morphological and molecular alterations, such as accumulation of lysosomes and misfolded proteins in the cytoplasm, changes in mitochondrial dynamics, modifications in chromatin structure and function, aberrations in gene expression, and the secretion of soluble components into the extracellular medium known as Senescence-Associated Secretory Phenotype (SASP), which can influence the microenvironment through paracrine signaling [2].

One of the most significant features of SCs is their ability to evade programmed cell death by developing and activating pathways that promote survival and resistance to apoptosis, known as SCAPs [3]. It has been proposed that this ability to evade apoptosis is the underlying reason for SCs accumulation in organs and tissues over time, thereby contributing to the onset and progression of various chronic degenerative diseases associated with aging [4]. The first SCAPs were identified by analyzing transcriptomic profiles of senescent and non-senescent human cells (preadipocytes and umbilical vein endothelial cells (HUVECs)). Among the genes identified are BCL-2/BCL-xL, PI3K/AKT, Serpins, ephrins, dependence receptors/tyrosine kinases, and hypoxia-inducible factor (HIF-1α) [5]. In subsequent studies, some other SCAPs have been identified [6, 7]. The discovery of SCAPs spurred the development of senotherapy, a pharmacological approach using senolytics - small molecules that selectively induce death of SCs by targeting SCAPs [8].

Although SCs can evade apoptosis by activating various SCAPs, this mechanism is complex and not yet fully understood [9, 10]. Several authors have proposed that the evasion of cell death is a consequence of the positive regulation of SCAPs, facilitated by the upregulation of survival and anti-apoptotic resistance genes and proteins [5, 7, 11, 12].

## Results and discussion

To explore this idea, we compared the expression levels of 39 genes involved in the SCAPs [5]. For this purpose, we used gene expression data from the GTEx project [13] from 33 different tissues obtained from young (20-39 years) and old (60-70 years) individuals. As illustrated in Figure 1 compares the distribution and mean expression levels quantified by transcripts per million (TPM) of the 39 genes. Statistical analysis revealed no significant differences in the mean expression of these 39 genes between young and old individuals despite assuming a higher accumulation of SCs in the tissues of aged individuals [14]. This result suggests that the anti-apoptotic resistance of SCs likely does not arise from a collective increase in the expression of SCAPs genes.

**Figure 1.**
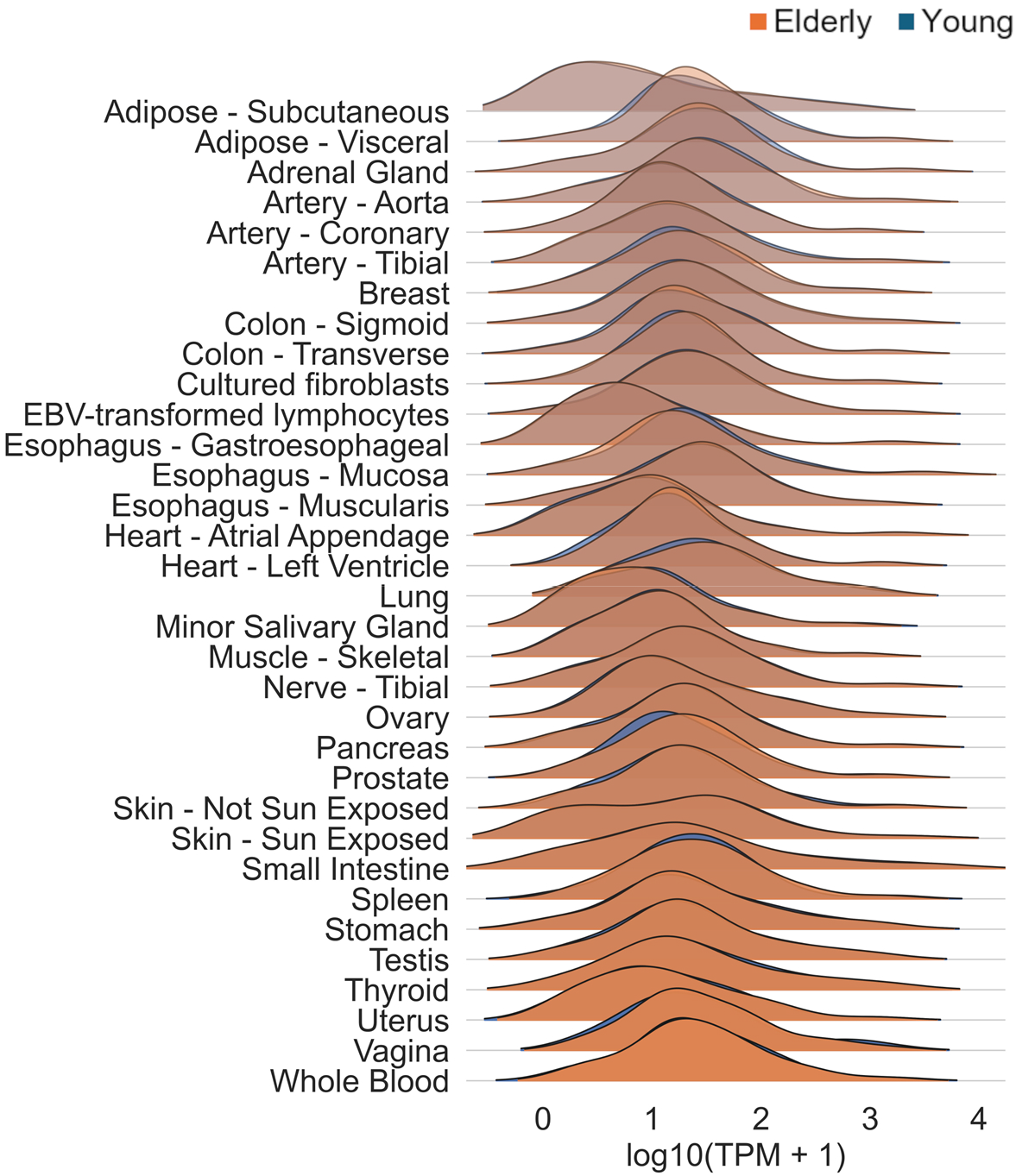
Analysis of the 39 genes involved in the SCAPs at the gene expression level. Comparison of the distribution and mean expression levels (transcripts per million) of the 39 genes collectively across 33 tissues from young (20-39 years) and old (60-70 years) individuals, (Student’s t-test, p < 0.05).

Furthermore, we analyzed the differential expression of the 39 genes during aging in each of the 33 tissues, as shown in Figure 2. It is observed that many these genes remain unchanged or without significant abrupt changes in their expression when comparing elderly to young individuals. Except for SERPINE1 and CDKN1A, which increased expression across multiple tissues, no gene showed a significant uniform increase in expression across all tissues. This suggests that the development and activation of SCAPs is tissue and/or cell type-dependent, which would explain why senolytics work in certain cell types and not others. Interestingly, a significant proportion of the genes decrease their expression levels in multiple tissues during aging. Additionally, some tissues did not show significant changes in the expression levels of these genes. The results suggest SCs’ SCAPs development/activation may not depend on differential expression of the comprising genes.

**Figure 2.**
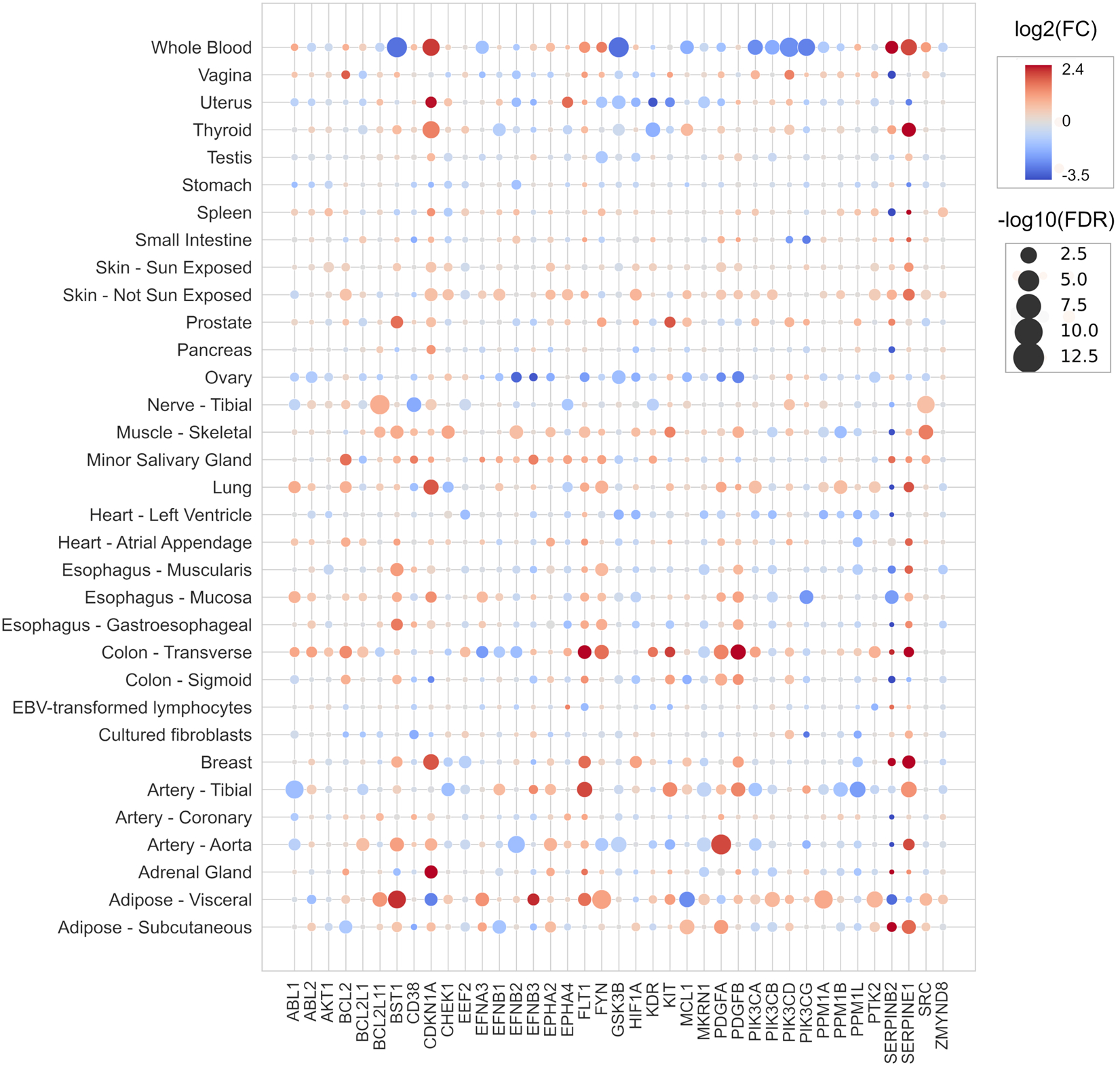
Analysis of the 39 genes involved in the SCAPs at the gene expression level. Differential expression of each of the 39 SCAPs genes across the 33 tissues, the log2(fold change (FC)) is represented by a color gradient indicating the increase or decrease in expression in tissues of aged individuals, and the size of the circles represents the -log10(False Discovery Rate (FDR)).

While alterations in the expression levels of a limited set of genes may influence certain cellular functions, they might not necessarily translate into phenotypic changes or concrete biological effects. This complexity arises because cellular functions result from a sophisticated network of interactions among various molecular components, and gene expression is one of several factors contributing to this intricate system. [15, 16]. For this reason, in this study, we propose using integrative methods that consider the collective behavior of the genes constituting the SCAPs. Consequently, we employed gene co-expression networks (GCNs) to examine the changes in the functional coordination of SCAPs genes [17]. This approach allows us to elucidate the intricate interplay between different genes within SCAPs and their role in age-related cellular changes.

To explore the correlation between the genes of the SCAPs, we constructed GCNs using mutual information (MI) as a metric of co-expression to quantify the statistical dependence between gene pairs [18]. As shown Figure 3, after inferring the GCNs in each of the 33 tissues in young and old individuals, we retained the 100 strongest interaction pairs from each network to quantify the strength of co-expression. Unlike gene expression levels, co-expression strength exhibited significant differences in most tissues according to age, being higher in older individuals. This finding is interesting because although aging has been reported to lead to a loss of gene coordination within GCNs [19].

**Figure 3.**
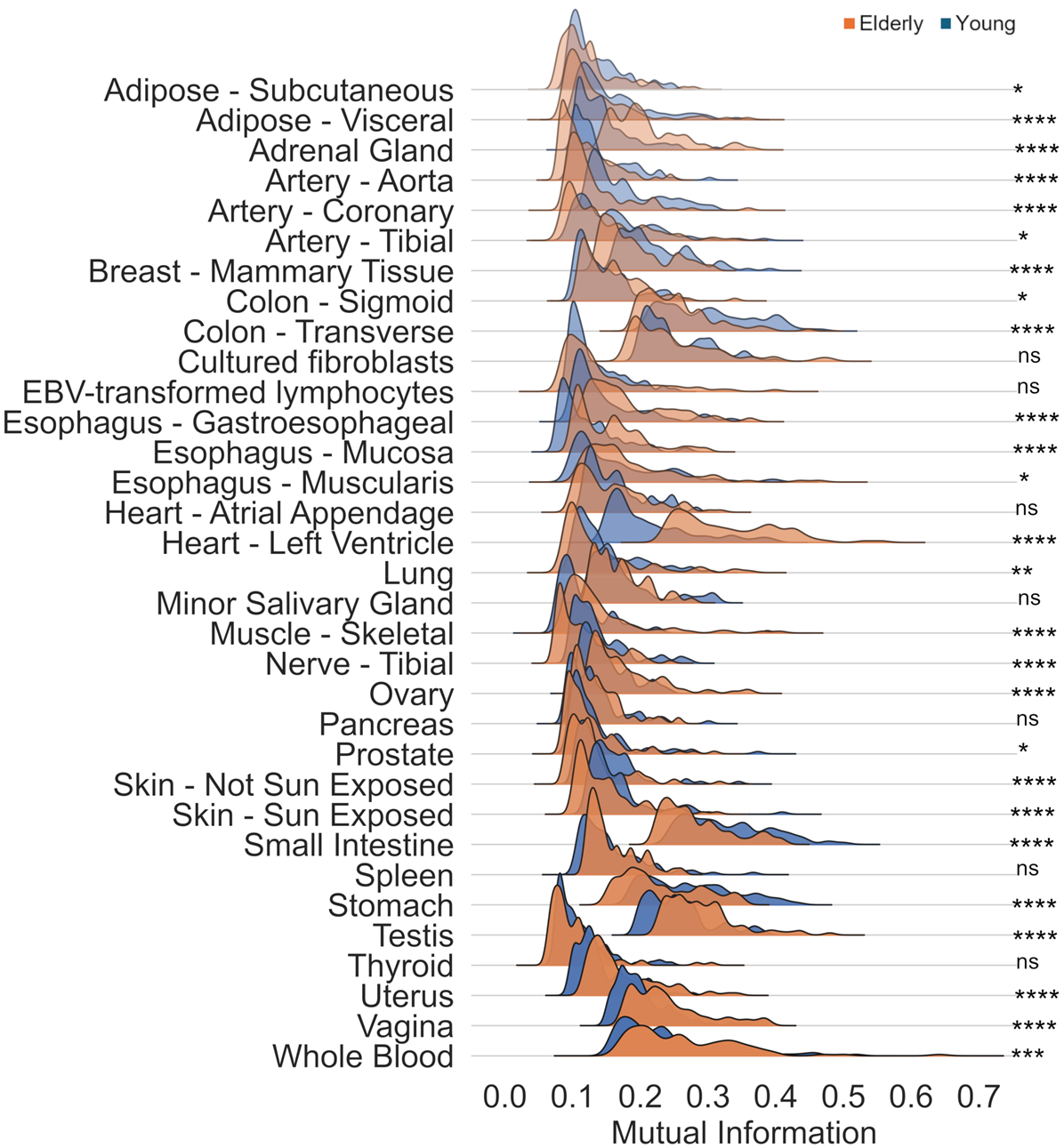
Analysis of gene co-expression networks (GCNs) of the 39 SCAPs genes. **A**. Distribution of mutual information values of the 100 strongest pairs of interactions from the networks formed by the 39 SCAPs genes in young and elderly individuals (Mann-Whitney U test). *p <= 0.01, **p < 0.001, ***p < 0.0001, ****p < 0.00001.

In addition to co-expression strength, another crucial factor influencing gene functional organization is the structure of GCNs, which refers to the specific gene wiring in each tissue. Co-expressed genes typically exhibit functional relationships or are regulated by the same transcriptional program [17]. Figure 4A illustrates the similarity of network wiring across different tissues in aged individuals. We utilized the Jaccard Index to measure interaction similarity (same genes connected by the same edges), and the results are depicted in a heatmap. Remarkably, the networks of the different tissues practically do not share interactions, implying a unique functional organization of SCAPs genes in each tissue. These disparities in gene functional coordination may significantly impact SCAPs development and activation, suggesting that each tissue harbors its distinct anti-apoptotic resistance mechanism.

**Figure 4.**
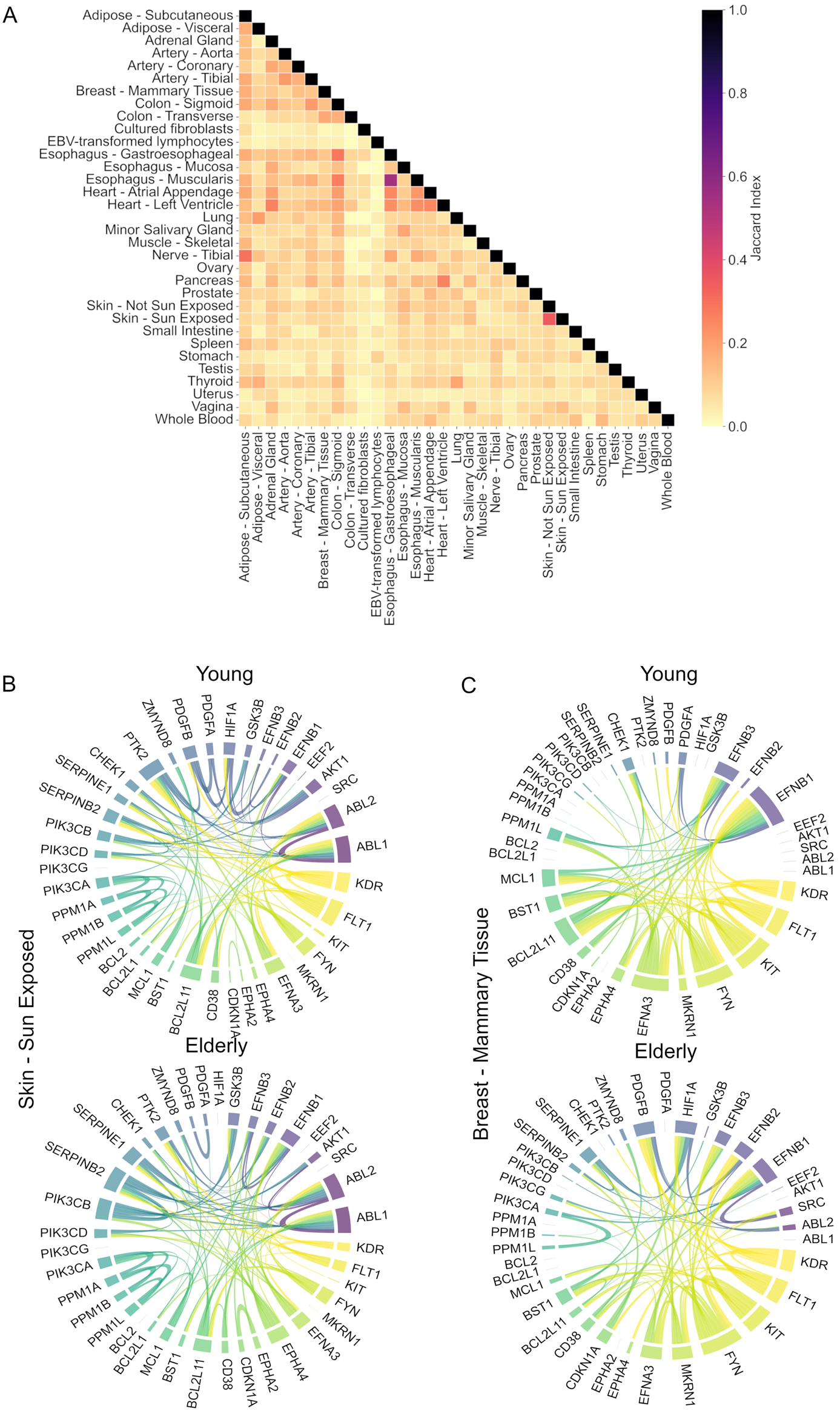
**A**. Connectivity of the GCNs, illustrating the number of shared interactions (genes connected by the same links) between the GCNs of different tissues in aged individuals. **B**. GCNs of Breast - Mammary Tissue. And **C**. GCNs of Skin - Sun Exposed Tissue. The GCNs and changes in their specific connections are depicted in tissues that gain, lose, and maintain their connectivity.

The differences in the co-expression patterns are evident between young and old individuals, as observed in panels B and C. These changes in connectivity are not uniform; they are heterogeneous. Some networks gain connections with age, such as the heart, where the atrial appendage GCN (Figure S1, Supplementary material) show denser connections in aged individuals. Conversely, networks like sun-exposed skin maintain consistent connections regardless of age, although there are notable changes in network wiring (Panel B). Additionally, some networks lose density with age but compensate by forming new connections, as observed in the breast mammary tissue (Panel C). These results suggest that the 39 SCAPs genes are not isolated entities but functionally interrelated and part of a complex, well-defined transcriptional program. Consequently, alterations in co-expression patterns due to aging could play a crucial role in the development and activation of SCAPs, indicating that SCs survival is an emergent property in response to the rewiring of GCNs.

## Conclusions

In this study, we have explored the behavior of some of the genes that make up the SCAPs, revealing their complexity and dynamism. Before this study, the prevailing notion proposed that the development and activation of SCAPs stem from the increased expression of specific genes within them; however, this conception could be reductionist. We suggest that the development and activation of SCAPs is a multifactorial process involving changes in expression levels and the strength and patterns of gene association. Moreover, these changes are likely tissue-specific and may vary based on the senescence-inducing stimulus and the senescence stage.

Gaining a deeper insight into how SCs evade apoptosis represents a significant advancement in the development of new senolytics. Such progress would mark a transition away from current agnostic targeting methods, which rely on hypothesis-driven and high-throughput phenotypic screening, towards more rational, targeted, and personalized design approaches. This shift holds promise for the discovery of senolytics that are not only more effective and potent but also more specific, thereby minimizing potential side effects.

Our findings offer new insights into the mechanisms by which senescent cells acquire resistance to apoptosis, potentially improving our understanding of this intricate process. However, interpreting these results with caution is of utmost importance due to their inherent limitations, as we used data from the tissues of elderly individuals. Although it is assumed that the accumulation of SCs at this stage of life is greater, this may only partially capture the true nature of pathogenic SCs.

## Methods

### Data Acquisition

The Genotype-Tissue Expression (GTEx) project provides gene expression data generated through bulk RNA-seq, specifically the GTEx V8 version offers data from 53 distinct tissues, collected from 714 donors without reported diseases whose ages ranged from 20 to 70 years [13]. Two datasets were obtained, the first of normalized expression data in transcripts per million (TPM) and the second of raw counts. In both original datasets, each sample included counts for a total of 56,200 genes.

For both datasets (normalized and raw counts) and for each of the 53 tissues, the dataset was divided into two different datasets of young (20 - 39 years) and old (60 - 70 years) individuals. Only tissues with more than 25 samples in each age group were selected, resulting in 33 tissues. The tissue and the number of samples per group used for the analyses are shown in Table 1.

**Table 1.** The tissue and the number of samples per group used for the analyses.

| **Tissue** | **Elderly** | **Young** |
| --- | --- | --- |
| Muscle - Skeletal | 292 | 132 |
| Whole Blood | 272 | 136 |
| Skin - Sun Exposed (Lower leg) | 260 | 116 |
| Artery - Tibial | 215 | 118 |
| Adipose - Subcutaneous | 237 | 110 |
| Thyroid | 234 | 98 |
| Nerve - Tibial | 229 | 104 |
| Esophagus - Mucosa | 166 | 107 |
| Skin - Not Sun Exposed (Suprapubic) | 226 | 97 |
| Esophagus - Muscularis | 143 | 110 |
| Lung | 212 | 73 |
| Breast - Mammary Tissue | 155 | 89 |
| Adipose - Visceral (Omentum) | 182 | 89 |
| Cells - Cultured fibroblasts | 162 | 83 |
| Artery - Aorta | 140 | 75 |
| Colon - Transverse | 105 | 90 |
| Heart - Atrial Appendage | 180 | 33 |
| Heart - Left Ventricle | 164 | 48 |
| Esophagus - Gastroesophageal Junction | 103 | 66 |
| Colon - Sigmoid | 135 | 75 |
| Stomach | 84 | 83 |
| Pancreas | 84 | 60 |
| Artery - Coronary | 82 | 26 |
| Adrenal Gland | 81 | 42 |
| Testis | 125 | 67 |
| Spleen | 54 | 52 |
| Prostate | 80 | 56 |
| Small Intestine - Terminal Ileum | 46 | 50 |
| Cells - EBV-transformed lymphocytes | 50 | 29 |
| Minor Salivary Gland | 52 | 30 |
| Ovary | 46 | 36 |
| Vagina | 42 | 28 |
| Uterus | 28 | 36 |

### Gene Expression Level Analysis

#### Collective Expression Levels

Using the TPM-normalized expression dataset, the mean expression values for each of the 39 SASP genes were extracted for each age group. Subsequently, a student’s t-test was performed for each tissue to compare the mean gene expression levels between the Young and Elderly groups, considering differences with a p-value less than 0.05 as significant. This test was carried out using the ttest_ind function from the scipy.stats library version 1.13.0. Finally, the visualization of the expression distributions was performed using the joyplot function from the joypy library version 0.2.6. For the visualization of gene expression distributions, the mean expression values of the genes in each tissue were transformed using the base 10 logarithm (log10). This preprocessing was carried out before generating the joyplot using the np.log10 function in the numpy 1.26.0 package.

#### Tissue-Specific Differential Expression

A non-parametric Mann-Whitney U test was implemented to compare gene expression levels between the Young and Elderly groups in each tissue. P-values were calculated for each gene and tissue using the mannwhitneyu function from the scipy.stats library. The obtained p-values were subjected to multiple testing correction using the Benjamini-Hochberg method to control the false discovery rate (FDR). This was done with the multipletests function from the statsmodels.stats.multitest library. To visualize the results, a scatterplot was generated where each point represents a gene in a specific tissue. The size and color of the points represent the adjusted p-value and the log2(fold change) in gene expression between the Young and Elderly groups, respectively. The color intensity indicates the magnitude of the change, while the point size indicates the statistical significance.

#### Gene Co-expression Network Analysis (GCNs)

##### Data Preprocessing

The raw counts of gene expression values for each tissue were preprocessed as follows: Initially, the mean of the counts for each gene across the samples was calculated. Genes with a mean below 10 were removed. Additionally, genes that presented zero-count values in more than 50% of the samples were discarded [20]. Subsequently, the dataset was quantile-normalized using the qnorm 0.8.1 library. The next step consisted of dividing the dataset into two groups: young and elderly. Protein-coding genes, long non-coding RNAs, microRNAs, pseudogenes, and other types of RNA species were retained.

#### Inference of GCNs

GCNs were inferred using mutual information (MI) as a measure of gene co-expression. Mutual information is a statistical measure that quantifies how much information one random variable, such as the expression of a gene, contains about another. The ARACNe (Algorithm for the Reconstruction of Gene Regulatory Networks) algorithm, one of the most popular methods for inferring GCNs, calculates the MI between two data series [21]. ARACNe was applied to both preprocessed datasets (young and elderly) to establish correlations between gene pairs in each tissue. The complete transcriptome was calculated for both datasets. To speed up the calculation, the multi-core C++ version without adaptive partitioning inference was employed [22]. This version is available at: https://github.com/josemaz/aracne-multicore.

#### Comparative Analysis of Co-expression Strength

The 100 strongest interactions from each network (young and elderly) in each tissue were retained to make them comparable. The MI values of the young and elderly networks for each tissue were compared using a non-parametric Mann-Whitney U test. The statistical test and the visualization of the distributions followed the same methodology reported above.

#### Analysis of Conserved Interactions across Different Tissues

Using the MI values of the complete transcriptome, the MI value corresponding to the 95th percentile of each network (young and elderly) in each tissue was obtained. These values were then used as a cutoff in each constructed GCN to obtain statistically significant networks. To investigate the interactions conserved across different tissues, an analysis was performed using the Jaccard index as a measure of similarity between the sets of gene interactions in each tissue. The Jaccard index is calculated as the size of the intersection of two sets divided by the size of their union, which allows quantifying the similarity between the network interactions. The result was then visualized using the heatmap function from the Seaborn 0.13 library.

The visualization of the tissue-specific wiring of the networks was constructed using the chord_diagram function from the mpl_chord_diagram 0.4.1 library.

## Author Contributions

S.O.: Conceptualization, formal analysis, figure design, writing and editing of the article. N.E.L.D: Funding, writing, and editing of the article. All authors approved the final version of the manuscript.

## Acknowledgments

We thank the Laboratorio de Supercómputo y Visualización en Paralelo (LSVP) of the Universidad Autónoma Metropolitana Campus Iztapalapa for the computing time granted on the ‘Yoltla’ supercomputer.

## Data availability

All data generated or analyzed during this study are public and included in this published article

## Code availability

The code used to support the analysis of this study can be found at https://github.com/Olascoaga/Survival-Networks

