## Supplementary material for "Towards a Rosetta Stone to Decipher the Survival Networks of Senescence": S1

Supplementary Figures

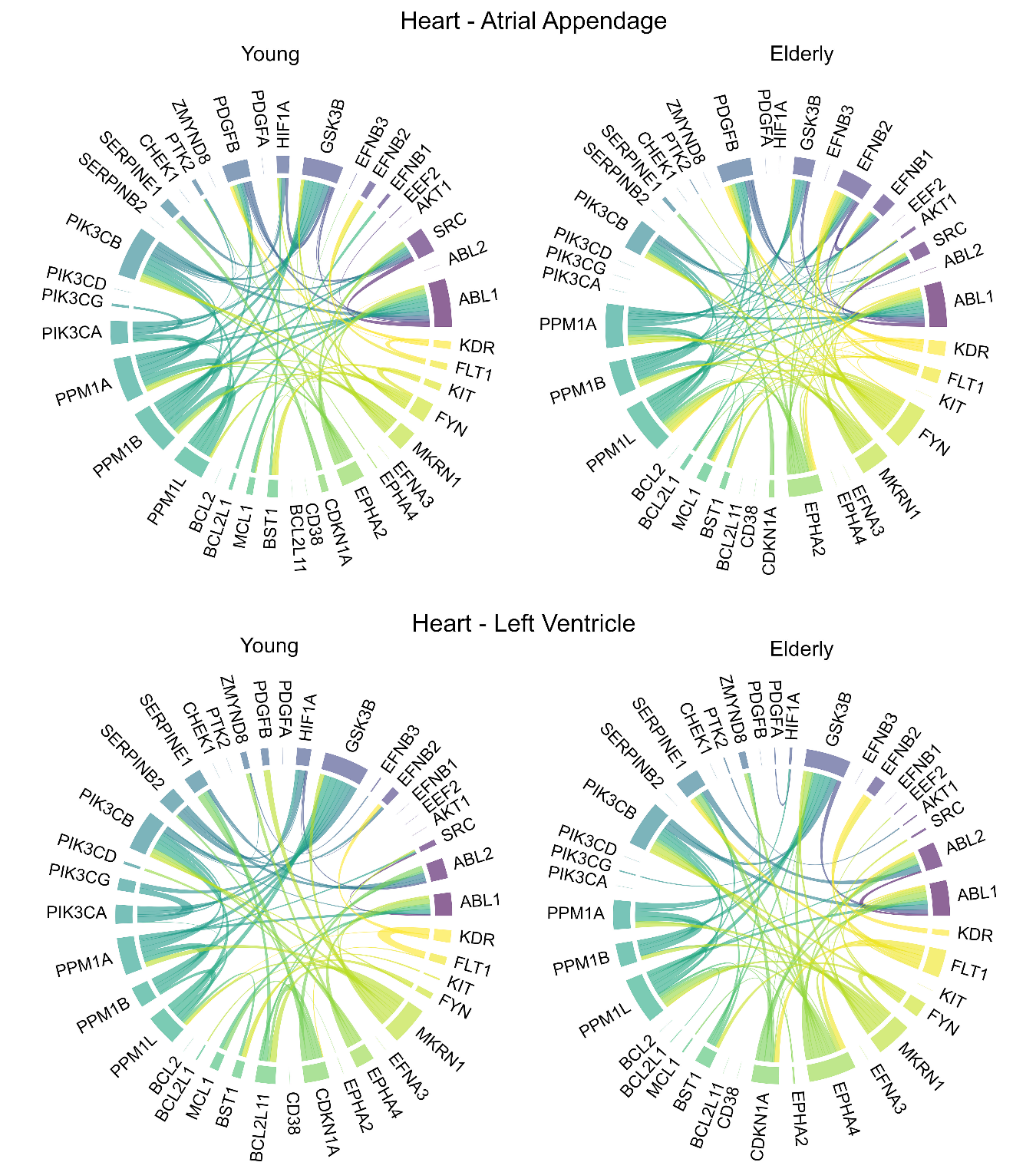

**Figure S1.** Gene co-expression networks of Heart - Atrial Appendage where an increase in network density (gain of connections) is observed and Heart - Left Ventricle where there is neither gain nor loss of connections but a change in co-expression patterns (rewiring of networks).
